# Applying meta-analysis to Genotype-Tissue Expression data from multiple tissues to identify eQTLs and increase the number of eGenes

**DOI:** 10.1101/100701

**Authors:** Dat Duong, Lisa Gai, Sagi Snir, Eun Yong Kang, Buhm Han, Jae Hoon Sul, Eleazar Eskin

## Abstract

During the last decade, with the advent of inexpensive microarray and RNA-seq technologies, there have been many expression quantitative trait loci (eQTL) studies for identifying genetic variants called eQTLs that regulate gene expression. Discovering eQTLs has been increasingly important as they may elucidate the functional consequence of non-coding variants identified from genome-wide association studies. Recently, several eQTL studies such as the Genotype-Tissue Expression (GTEx) consortium have made a great effort to obtain gene expression from multiple tissues. One advantage of these multi-tissue eQTL datasets is that they may allow one to identify more eQTLs by combining information across multiple tissues. Although a few methods have been proposed for multi-tissue eQTL studies, they are often computationally intensive and may not achieve optimal power because they do not consider a biological insight that a genetic variant regulates gene expression similarly in related tissues. In this paper, we propose an efficient meta-analysis approach for identifying eQTLs from large multi-tissue eQTL datasets. We name our method RECOV because it uses a random effects (RE) meta-analysis with an explicit covariance (COV) term to model the correlation of effect that eQTLs have across tissues. Our approach is faster than the previous approaches and properly controls the false-positive rate. We apply our approach to the real multi-tissue eQTL dataset from GTEx that contains 44 tissues, and show that our approach detects more eQTLs and eGenes than previous approaches.

## 1 Introduction

Expression quantitative trait loci (eQTL) studies aim to discover genetic variants called eQTLs that influence gene expression by collecting information on both gene expression and genetic variants using microarray or sequencing technologies (The GTEx Consortium, 2015). Such variants may be identified by testing correlation between a genotype of a variant and expression levels of a gene. Recent genome-wide association studies (GWAS) have discovered that many disease causing variants are eQTLs, and these eQTLs may play an important role in regulation of genes and complex traits (Albert, 2016; Liu <u>et al.,</u> 2016; Nieuwenhuis <u>et al.,</u> 2016). Disease-causing variants may affect higher level traits through gene expression, and determining whether or not a variant is an eQTL would help us understand its functional impact on the trait (Guauque-Olarte <u>et al.,</u> 2015; Choi <u>et al.,</u> 2016; Richard <u>et al.,</u> 2016).

Recently, improvements and cost reduction in microarray and RNA-seq technologies have allowed eQTL studies to collect gene expression from multiple tissues for a better understanding of the functional mechanisms of variants. For example, the Genotype-Tissue Expression (GTEx) consortium has performed RNA-seq in more than 40 tissues from hundreds of individuals (The GTEx Consortium, 2015). These multi-tissue eQTL datasets provide not only biological insights into the effect of genetic variants in multiple tissues but also greater statistical power to detect eQTLs shared across tissues. If an eQTL has small effect in several tissues, one may combine information from those tissues and boost power to detect its effect. There are a few methods designed to detect eQTLs from multiple tissues such as Meta-Tissue and eQTLBma (Flutre <u>et al.,</u> 2013; Sul <u>et al.,</u> 2013). Meta-Tissue uses both linear mixed models (LMM) and meta-analysis to aggregate statistics from multiple tissues, and eQTLBma is a Bayesian model that combines data from several tissues where a variant is allowed to have unique effects in different tissues (Flutre <u>et al.,</u> 2013; Sul <u>et al.,</u> 2013). These methods take into account the fact that an eQTL may have different effect across tissues, a phenomenon known as heterogeneity. They are shown to identify more eQTLs than a traditional "tissue-by-tissue" (TBT) approach that examines each tissue separately.

These methods, however, do have several limitations that may lower their statistical power and applicability to large eQTL datasets. One major limitation is that they are computationally intensive and cannot be applied to multi-tissue datasets containing many tissues (e.g. > 20 tissues). Meta-Tissue involves estimating variance components for every pair of a variant and gene expression in its LMM framework, which is a computationally heavy procedure when there are thousands of samples in all tissues (Sul <u>et al.,</u> 2013). eQTLBma uses a Bayesian framework that considers all possible combinations of tissues in which an eQTL has effect, and this corresponds to 2^*T*^ configurations where *T* is the number of tissues (Flutre <u>et al.,</u> 2013). This approach is computationally infeasible when *T* is 44 such as in the GTEx data. Hence, the GTEx consortium used a meta-analysis software called Metasoft, which is equivalent to Meta-Tissue without the LMM framework (Han and Eskin, 2011, 2012). Metasoft extends the random-effects (RE) model of meta-analysis to attain better statistical power; the extended RE model is named RE2 (Han and Eskin, 2011,2012). This approach, however, may not be an optimal choice as it assumes tissues do not share samples from the same individual. This assumption is often violated in multi-tissue eQTL datasets because samples from different tissues are often collected from the same donor.

Another limitation in these previous methods, especially of Meta-Tissue, is their assumption that a genetic variant has independent effects in multiple tissues. This assumption is, however, often invalid in multi-tissue eQTL studies because a genetic variant tends to have similar effect in related tissues (The GTEx Consortium, 2015). For example, a variant may similarly regulate expression of a certain gene in several adjacent brain tissues, and this means that it may be an eQTL in all those tissues. Ignoring this correlation of effect of eQTLs in multiple tissues may reduce the statistical power to detect eQTLs.

In this paper, we propose a novel meta-analysis approach for multi-tissue eQTL studies that addresses all these limitations. Our method is based on RE2 meta-analysis and explicitly models correlation of effect that a genetic variant has in multiple tissues. We name our method RECOV because it uses a random effects (RE) meta-analysis with an explicit covariance (COV) term to model the correlation of effect that eQTLs have across tissues. Our method is markedly more efficient than previous approaches and can detect eQTLs from 44 tissues in GTEx data. In addition to identifying eQTLs, our method can identify eGenes; these are genes that contain at least one eQTL. Many eQTL studies are interested in identifying eGenes because they are more interpretable and can be analyzed as a network or a pathway. We use simulations to show that our approach has correct false positive rates even when it is applied to eQTL datasets with many tissues. We then apply our approach to real multi-tissue eQTL data from GTEx and show that our approach detects more eGenes than the previous RE2 and TBT approaches.

## 2 Method

We begin by introducing the notations that are used in this paper. We use the notation *x* ∈ ℝ^*n*^ to specify a vector *x* with dimension *n*, and *Z* ∈ ℝ^*n*×*m*^ to specify a matrix *Z* with dimension *n* × *m*. *x_i_* denotes the *i*^th^ element in *x*, and likewise, we use *Z_ij_* to specify the *ij* entry in *Z*. We denote an item *k* in the set *K* by *k* ∈ *K*, and a set {*a*_1_…*a_K_*} indexed by *k* by using {*a_k_*}_*k*∈*K*_, and the subscript *k* ∈ *K* is omitted whenever the context is clear. The size of the set *K* is denoted as |*K*|.

### 2.1 Detecting one eGene via an eQTL study

#### 2.1.1 eQTL study in one tissue

We begin with an eQTL study in one tissue *t*. An eQTL study finds every eQTL associated with the expression level of a specific gene *g*. To do this, the study tests each variant *v* in the set *V* against the expression of *g* in a sequential fashion. To set up the problem, suppose we represent the gene expression for *m* individuals in tissue *t* as a vector *q* ∈ ℝ^m^, and we want to find the effect of variant *v* on *g*. Let *S* ∈ ℝ^*m*^ be the standardized genotypes of this *v*. eQTL study assumes the following model

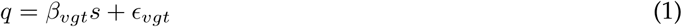

where *∈* ∈ ℝ^*m*^ is the sampling errors 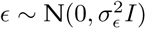, and *β_vgt_* ∈ ℝ is the true effect size of the variant *v* on *g* in tissue *t* (Darnell <u>et al.,</u> 2012; Eskin, 2015; Hormozdiari <u>et al.,</u> 2015). The estimate *b_vgt_* of the true value *β_vgt_* can be computed using the basic least squares equation 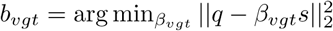. This solution is

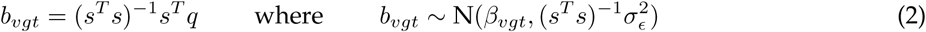

(Abraham and Ledolter, 2006). By using Eq. 2 and writing the null hypothesis H_0_: *β_vgt_* = 0, one can do a hypothesis test to assert if *v* has an effect toward *g*. To do this test, compute the estimate 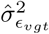 of 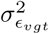 by

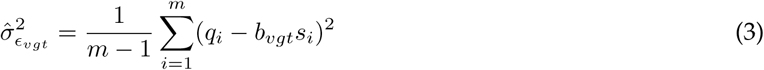

and estimate the variance *d_vgt_* of *b_vgt_* by

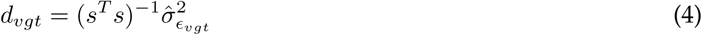

then compute the p-value *p_vgt_* = p-value(*b_vgt_*) (Abraham and Ledolter, 2006; Eskin, 2015). If *p_vgt_* is less than some significance level, then we reject H_0_: *β_vgt_* = 0, and conclude that *v* is an eQTL of *g* in tissue *t*. When many variants are tested, multiple testing correction is needed due to linkage disequilibrium (LD); for example, one can apply Bonferroni correction by using the threshold *α*/|*V*| where *α* is the significance level for the whole family of tests. There exist other methods that can handle LD in the set *V* better than the Bonferroni correction (Conneely and Boehnke, 2007; Joo <u>et al.,</u> 2014; Hormozdiari <u>et al.,</u> 2015; Joo <u>et al.,</u> 2016).

#### 2.1.2 Using an eQTL study in one tissue to discover one eGene

Because an eQTL study tests each variant *v* ∈ *V* against gene *g* in a tissue *t*, from one single eQTL study, we have a set of p-values {*p_vgt_*}_*v*∈*V*_. The minimum *p_gt_* = min_*v*∈*V*_ {*p_vgt_*} is defined to be the observed eGene statistic at gene *g* in tissue *t* (The GTEx Consortium, 2015). Define *α_pgt_* = p-value(*p_gt_*) to be the eGene p-value (The GTEx Consortium, 2015). The eGene p-value depends on two important factors: the number of variants |*V*|, and the LD of the variants. In practice, *α_pgt_* is computed by doing a permutation test (Sul <u>et al.,</u> 2015; Duong <u>et al.,</u> 2016). In brief, in the *k*^th^ permutation, one would permute the gene expressions among the individuals, and compute a new 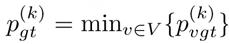. *α_pgt_* is the ranking of the observed *p_gt_* with respect to the density created from many 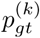. One can then conclude that *g* is an eGene in tissue *t* if its eGene p-value *α_pgt_* is less than some desired threshold.

### 2.2 Tissue-by-tissue analysis to find one or many eGenes

When there are genotype-tissue expression data from many tissues, the tissue-by-tissue (TBT) analysis is the standard method to find eGenes (Sul <u>et al.,</u> 2013; The GTEx Consortium, 2015). TBT analysis examines each tissue individually. TBT tests whether or not the gene has at least one eQTL in each tissue. Suppose the gene is expressed in *T* tissues. TBT performs *T* eQLT studies (one test in each tissue). The null hypothesis is that the gene is not eGene in any tissue. This hypothesis is equivalent to saying that no eQTL is found for this gene in any tissue.

Three layers of multiple testing correction are required since TBT performs one test per gene in each tissue. The first layer of multiple testing correction is applied within a tissue and corrects for LD of the variants tested against the gene. This correction can be done by using the permutation test to compute the eGene p-value for the gene in the tissue (Sul <u>et al.,</u> 2013, 2015; The GTEx Consortium, 2015; Duong <u>et al.,</u> 2016).

The second layer of multiple testing correction adjusts for the fact that we may test more than one gene within one tissue. For example, the GTEx pilot study tested thousand of genes within one tissue, and then transformed eGene p-values into eGene q-values to control for this multiple testing (Dabney <u>et al.,</u> 2010; The GTEx Consortium, 2015). This second layer of multiple testing correction is not needed if only one gene is tested in each tissue.

The third layer of multiple testing correction takes into account the fact that one gene is tested T number of times (one time for each tissue) (Sul <u>et al.,</u> 2013). In this layer, we apply Bonferroni correction so that the false-positive threshold for any eGene q-value in each tissue is *α/T*, where *α* is 5% for example.

### 2.3 Meta-analysis models for combining eQTL studies across tissues

We motivate the application of meta-analysis for combining eQTL studies across tissues. An eQTL is defined not only with respect to a gene, but also with respect to the tissue in which the gene expression is measured. eQTL studies of the same gene have been analyzed separately at the tissues level (Sul <u>et al.,</u> 2013). We can better detect the effect of a variant on the gene by combining eQTL results across many tissues and modeling the relatedness of the effect sizes of one variant among the tissues.

It is important to emphasize that, when using meta-analysis to find many eGenes, one would need only two layers of multiple testing correction. The first layer is applied within a gene to correct for LD because one tests many variants against the gene. The second layer is applied at the gene level because there is usually more than one gene being tested.

We define the notations to be used later. Suppose we have *T* eQTL studies (one study per tissue) that test the association of a variant *v* at a gene *g*. As shown above, denote the effect of this variant in the study (i.e. tissue) *t* as *b_vgt_*, where *b_vgt_* is computed using Eq. 2. Denote the variance of *b_vgt_* in the study *t* as *d_vgt_* where *d_vgt_* is computed using Eq. 4. Let *b_vg_* ∈ ℝ^T^ contains the effects in these *T* studies, so that *b_vg_* = [*b*_*vg*1_ *b*_2*vg*_…*b_vgt_*]^┬^. Let *D_vg_* = **diag**(*d*_*vg*1_…*d_vgT_*).

#### 2.3.1 Random effects (RE) and the RE2 model

The maximum likelihood procedure in RE model assumes that *b_vg_* has the form (Thompson and Sharp, 1997; Han and Eskin, 2011)

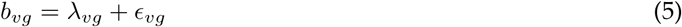

The random sampling errors *∈_vg_* are estimated from the data and assumed to be *∈_vg_* ∼ N(0, *D_vg_*). λ_*vg*_ ∈ ℝ^*T*^ in the RE model is a random variable, that is 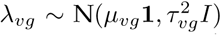 with 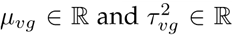. The effect λ_*vg*_ is thus known as the random effect. *μ_vg_* is the common true underlying effect that all the studies inherit. The term 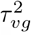 is the heterogeneity among the effects of the variant in *T* tissues.

Clearly, *b_vg_* comes from the distribution

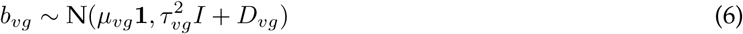

The traditional RE model assumes that if the effect of the variant does not exist in any tissue, then *μ_vg_* = 0. However, it is shown that this traditional null hypothesis does not yield optimal statistical power in detecting eQTLs (Han and Eskin, 2011, 2012). For this reason, the RE2 model assumes a different null hypothesis. If the effect of the variant does not exist in any tissue, then *μ_vg_* = 0 and 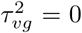. The fact that 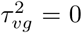 is a result of *μ_vg_* = 0, because when the effect does not exist, then its variance must not exist (Han and Eskin, 2011, 2012; Kang <u>et al.,</u> 2014). We will compare our method against the RE2 model.

The null hypothesis H_0_ in RE2 is

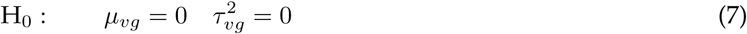

The likelihood ratio test becomes

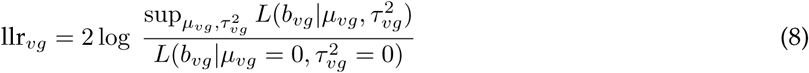

The function *L* denotes the likelihood function of the random variable *b_vg_*. The numerator 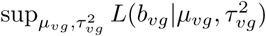 has to be estimated using a grid search or numerical methods based on the restricted maximum likelihood (REML) (Corbeil and Searle, 1976; Fraley and Burns, 1995; Gilmour <u>et al.,</u> 1995). There are also other derivative free methods. Here, we apply the Nelder-Mead method, which is a heuristic derivative free search method.

In finding the supremum, one implicitly enforces 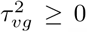. Due to this restricted parameter space, the asymptotic density of the likelihood ratio is an average of a 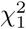 and 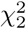 (Self and Liang, 1987; Han and Eskin, 2011). To find the p-value of this likelihood ratio when *T* is large, one can use this asymptotic density.

Otherwise, an alternate choice to compute the likelihood ratio p-value is to create a density of likelihood ratios under the null hypothesis and rank the observed likelihood ratio with respect to this density. One can make this null density by sampling many instances of *b_vg_* using Eq. 6 with 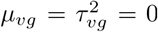, and computing their corresponding llr_*vg*_. If the p-value of llr_*vg*_ is significant, then *v* is an eQTL with respect to *g* in at least one tissue. Because we have 44 tissues in the GTEx data, we will use the asymptotic distribution of the likelihood ratio.

#### 2.3.2 RECOV: Random effects (RE) model with a covariance (COV) term

Here we present an extension to the RE model. We first discuss the covariance term. Eq. 5 of the RE model assumes 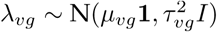 so that the effects of variant *v* toward gene *g* are independent across the tissues. However, tissues from the same body part are similar; in fact, many eQTLs are found to be consistent among many tissues (Flutre <u>et al.,</u> 2013). From this observation, we must acknowledge that λ_*vg*_ ˜ N(*μ_vg_* **1**, ∑_*vg*_) where ∑_*vg*_ is not diagonal. The term ∑_*vg*_ ∈ ℝ^*T*×*T*^ models the covariance of effect sizes of *v* among tissues conditioned on the gene *g*. In practice, ∑_*vg*_ must be estimated. Here we assume ∑_*vg*_ ≈ *c_vg_U_vg_* where *c_vg_* ≥ 0 is a scaling constant and is to be optimized jointly with the mean of regression coefficient *μ_vg_*.

In this paper, we compute the *U_vg_* at each variant-gene pair as follows. Denote *B_g_* = [*b*_1*g*_ *b*_2*g*_…*b*_|*V*|*g*_] so that *B_g_* ∈ ℝ^*T*×|*V*|^. Thus, a column in *B_g_* contains the effects of a variant in 44 tissues. To avoid reusing the data when testing a single SNP, we remove its effects in 44 tissues when estimating its covariance term. To do this, we divide all cis-variants of *g* into 10 separate segments according to their physical locations on the chromosome, and use the 9 segments that do not contain *v* to compute *U_vg_*. In particular, denote *B_−vg_* as the matrix *B_g_* without the effect sizes of the variants that belong to the same segment as *v*. *U_vg_* can be estimated as 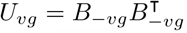 (after proper scaling applied to *B_−vg_*). This computation is similar to how one would compute a kinship matrix using the genotype matrix (Eskin, 2015). In this scheme, we also hope that the variants in strong LD with *v* are removed, so that there are fewer vectors in *B_−vg_* that resembles *b_vg_* when computing *U_vg_*.

Now, we are ready to introduce this covariance term *U_vg_* to the RE model. We extend the RE model so that when testing a variant *v* against gene *g*, we have

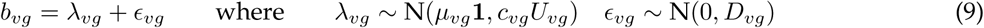

Like before, the matrix *D_vg_* is assumed to be known. Clearly,

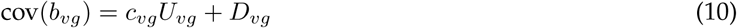

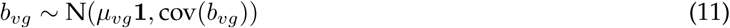

The null hypothesis that *v* does not effect *g* in all *T* tissues is

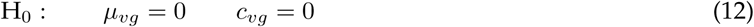

The alternative hypothesis implies that *v* has an effect in at least one of the *T* tissues.

Under this setting, the log likelihood ratio to test the hypothesis becomes

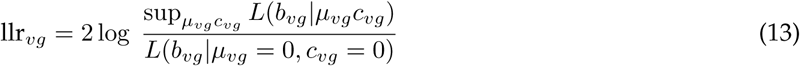

Like in the RE model, in finding the supremum in the alternative, one enforces *c_vg_* ≥ 0. Due to this restricted parameter space, the asymptotic density of the likelihood ratio is an average of a 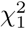 and 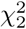. Alternatively, one can compute the empirical p-value of this likelihood ratio with a permutation test. In any case, if the p-value of the likelihood ratio is significant, then *v* is an eQTL with respect to *g* in at least one tissue.

### 2.4 Using meta-analysis of eQTL in many tissues to identify eGenes

In practice, a set of variants *V* is tested against *g*. Here we describe how one can combine the meta-analysis result at each variant *v* ∈ *V* to determine if *g* is an eGene.

Define *p_vg_* = p-value(llr_*vg*_) so that from many variants, we have the set of p-values {*p_vg_*}_*v*∈*V*_. The observed statistic at gene *g* is *p_g_* = min_*v*∈*V*_ {*p_vg_*}. To determine if *p_g_* is significant, one needs to compute its eGene p-value denoted as *α_p_g__* (The GTEx Consortium, 2015). To control for multiple testing when LD of the variants exists, one can compute *α_p_g__* using a permutation test (The GTEx Consortium, 2015; Sul <u>et al.,</u> 2015; Duong <u>et al.,</u> 2016). The permutation test creates a distribution of the observed *p_g_* under the null hypothesis. Using this distribution, one can compute the eGene p-value *α_p_g__* of *p_g_*.

This permutation test can be done as follows. Let *K* be the number of permutations. In the *k*^th^ permutation, permute the gene expression of *g* among the individuals in each of the *T* tissues so that there are *T* permuted data sets. This permutation reflects the hypothesis that the gene is not an eGene in any tissue. Next, redo the meta-analysis at each variant *v* ∈ *V* so that a new 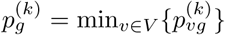 is computed. *α_pg_* is the fraction of times the observed *p_g_* is less than 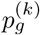. The gene *g* is an eGene in at least one tissue if its eGene p-value *α_p_g__* is below some threshold *α*.

In the pilot GTEx analysis, a set of genes *G* is being tested at once, so that one has a set *α_G_* = {*α_pg_*}_*g*∈*G*_. To control for the family wise error rate, one can apply Bonferroni correction to get the threshold *α*/|*G*|. Any gene *g* ∈ *G* with *α_pg_* < *α*/|*G*| is an egene in at least one of the *T* tissues.

#### 2.4.1 Estimating eGene p-value

The permutation test above must be performed at every pair of variant *v* ∈ *V* and gene *g* ∈ *G* in a tissue *t*. The entire permutation test requires *K*|*V*||*G*|*T*| permuted datasets, which is time consuming. Here, we introduce an alternate method to estimate the eGene p-value. The permutation test in essence estimates a function *f* that maps a test statistic to its p-value. There is evidence that the correlation of test statistics at two variants is equal to their LD (Han <u>et al.,</u> 2009). This holds true when the test statistics are effect sizes (Han <u>et al.,</u> 2009).

In our meta-analysis, to properly estimate the eGene p-value *α_p_g__*, we must consider the effect of LD in the set of variants *V* on the observed statistic at each variant *v*. At each variant, it does not matter whether we treat its llr_*vg*_ or its *p_vg_* as the observed statistic, because the likelihood ratio and its p-value are two equivalent entities for two reasons. First, the likelihood ratio of each variant *v* has the same distribution and degree of freedom. Second, the p-value function is 1-to-1 and strictly monotone. Thus, having a null density for the max_*v*∊*V*_{llr_*vg*_} is equivalent to having a null density for the minimum *p_g_* = min_*v*∈*V*_{*p_vg_*}.

We empirically find that on average the correlation of the likelihood ratios at any two variants is roughly equal to their LD; that is, on average cor(llr_*ug*_, llr_*vg*_) ≈ LD(*u*, *v*) for any variant *u*, *v* ∈ *V* (Figure 1). For this reason, any function *f* that accounts for LD and maps an observed test statistic of a gene into an eGene p-value would be applicable in our case. We can use such a function to convert the observed test statistics *p_g_* at the gene *g* to eGene p-value *α_p_g__* without doing the permutation test. Each gene g has its own LD and requires its own function f, because the cis-variants of each gene are non-identical.

We apply MVN-EGENE to estimate the function *f* for each gene. MVN-EGENE is a software that tests if a gene is an eGene in one tissue. To compute the function *f* at a gene, we apply MVN-EGENE at that gene in a tissue (Sul <u>et al.,</u> 2015). We assume that the LD does not change much between tissues thus it does not matter much about the particular chosen tissue, as long as it has many samples.

**Figure 1:**
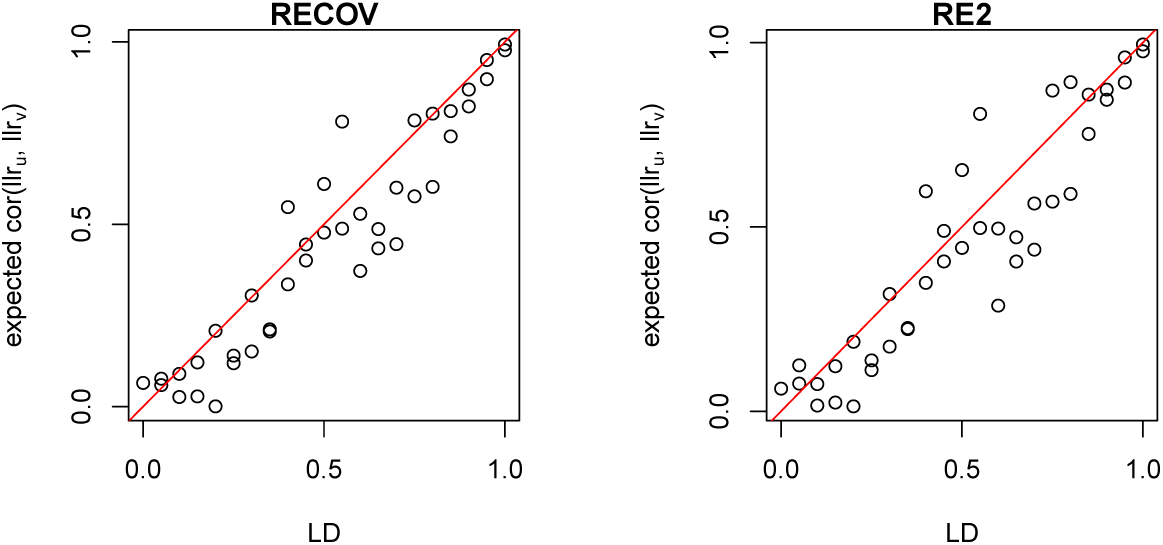
Relationship between likelihood ratios of a pair of variants and their LD. A SNP may be a cis-SNP for multiple genes and therefore tested as an eQTL for multiple genes. Denote cor(*llr_u_*, *llr_v_*) as the correlation between the likelihood ratio statistics of variants *u* and *v* in all genes where they are both cis-SNPs. Using cis-SNPs from the gene ENGS00000204219.5, we randomly selected pairs of SNPs that appeared in at least two genes together. The pairs were chosen such that 20 pairs were selected from LD bins of of width 0.05. and computed the correlation of their likelihod ratios in any other genes where they appear as a cis-SNP. We used both RECOV and RE2 meta-analysis to compute the likelihod ratios at each variant. We then find the mean cor(llr_*u*_, llr_*v*_) in each LD bin, using either RECOV (left) or RE2 (right) likelihood ratios. Plots show correlations after taking absolute value. The identity line is shown in red.

In MVN-EGENE, the test statistic for a gene is the most significant effect size taken over all cis-variants. The p-value of this test statistic depends on the LD of the cis-variants. Instead of doing a permutation test to compute this p-value, MVN-EGENE simulates data under the null hypothesis using a multivariate normal distribution. In brief, in one simulation, MVN-EGENE samples the effect sizes of the cis-variants of a gene in a tissue using zero as the mean effect and LD as the covariance matrix. In this simulation, the most significant effect among these effect sizes is taken to be the test statistic at the gene. After many simulations, one can create a null distribution for the observed test statistic. One can easily convert an effect size into a p-value using a normal distribution. By having a null density of the most significant effect size taken over all the variants, one also has the null density of the minimum p-value taken over all the variants. This null density of the minimum p-value in MVN-EGENE properly handles LD at the gene. Here, we use this distribution of minimum p-values as our null density to convert the observed minimum likelihood ratio p-value *p_g_* to its eGene p-value *α_p_g__* in both RECOV and RE2.

#### 2.4.2 Estimating genomic control

In the GTEx data set and other multi-tissue gene expression datasets, the same individual may provide samples for many tissues (Figure 2). LD in variants and shared individuals in tissues are both known to inflate the false positive rate in a meta-analysis (Sul <u>et al.,</u> 2013; Han <u>et al.,</u> 2016). Section 2.4.1 above describes how RECOV and RE2 handle LD in the variants. We now describe how we use a genomic control (GC) factor to remove the effect of shared individuals in the tissues from the meta-analysis results.

**Figure 2:**
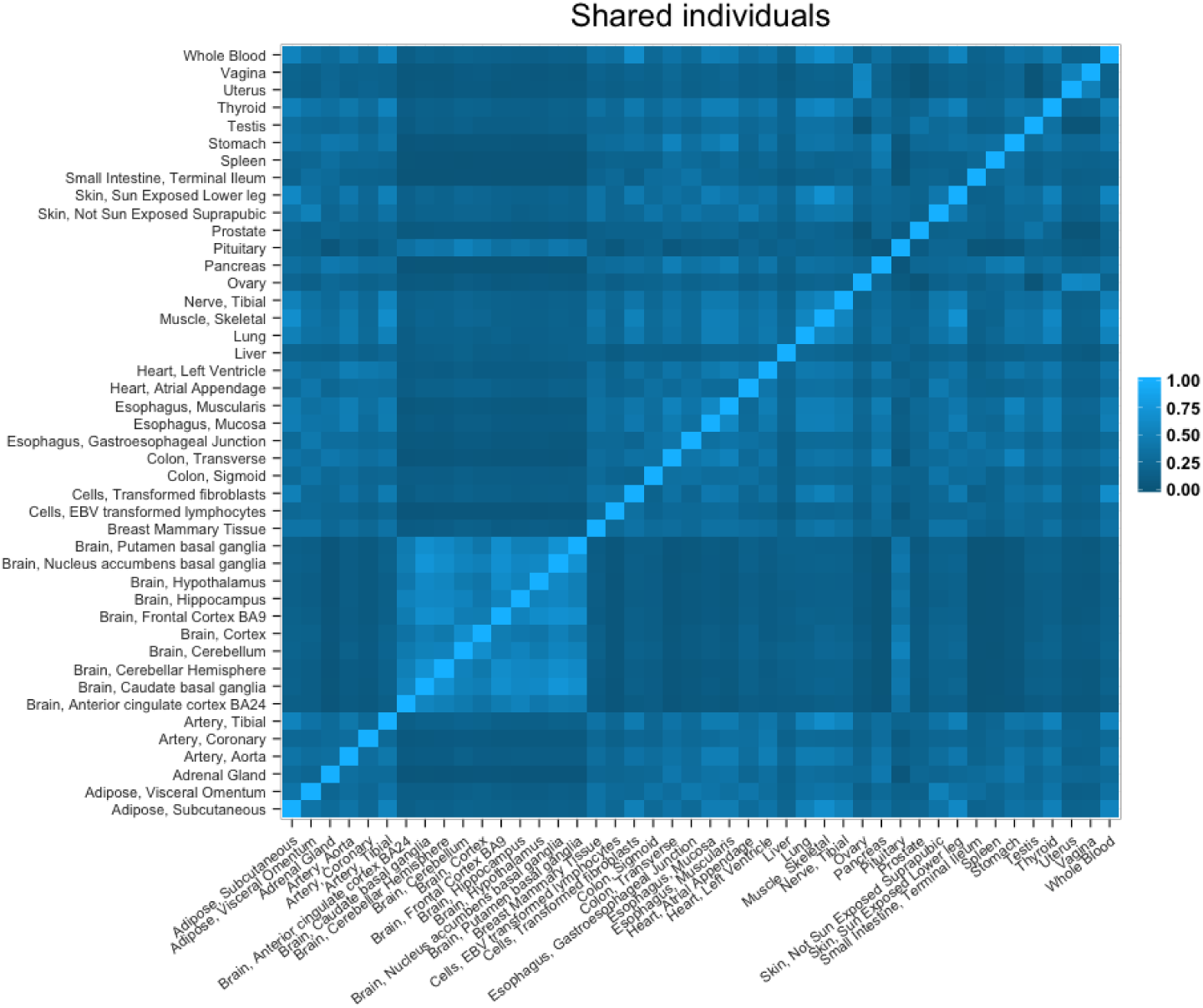
Shared individuals among the 44 tissues in the GTEx dataset. Degree of overlap of individuals between two tissues is measured using the Jacquard index.

Before testing whether the RECOV outcome is affected by the fact that tissues share individuals, we test if RECOV inflates the false positive rate when the data is absolutely free of any statistical association. When there is not any statistical association in the data, any sort of alternative hypothesis must be rejected more often than the null hypothesis. We simulate data to demonstrate that RECOV does not inflate false-positive rate in this case. These simulated datasets are free from signal due to LD, shared individuals in tissues, or correlated expressions of the same gene (or between genes) across tissues. In each simulated dataset, the number of individuals per tissue is taken from the GTEx data but we do not let tissues share individuals. We generate 1,000 SNPs at various minor allele frequency (MAF) without LD, and a random gene expression in each tissue. Between any two tissues, we do not make the expression of the same gene to be correlated. We compute the p-value of the likelihood ratio at each SNP using both RECOV and RE2 model. We repeat this simulation 1,000 times to obtain 1,000,000 p-values each for RECOV and RE2. The histograms of these p-values in both RECOV and RE2 indicate that the null hypothesis is more favored than the alternative hypothesis (Figure 3A,3B).

**Figure 3:**
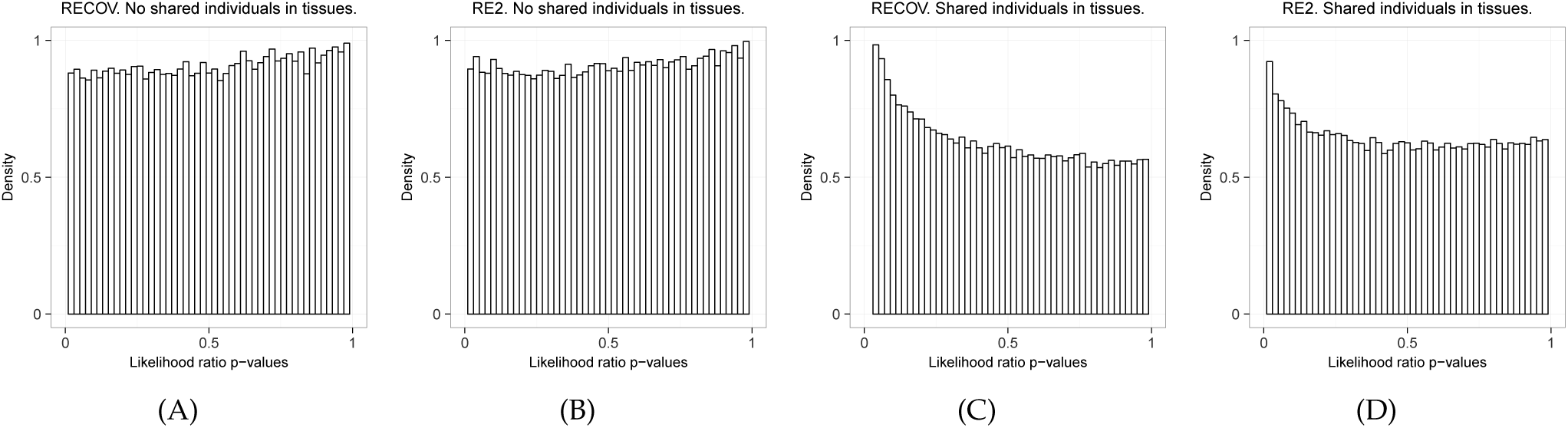
(A) RECOV and (B) RE2 applied to datasets where the tissues do not share individuals. (C) RECOV and (D) RE2 applied to datasets where the tissues share individuals.

To measure the effect strictly caused by shared individuals, we simulate datasets as above, but we allow tissues to share individuals. The number of people shared between pairs of tissues is taken from the GTEx data. As shown above, in each simulation we compute the likelihood ratio p-values at 1,000 SNPs, and repeat the simulation 1,000 times to obtain 1,000,000 p-values. We observe that these p-values shift toward 0 when the tissues share samples that are from the same individuals (Figure 3C,3D). In this case, we estimate the RECOV and RE2 GC factor to be 1.2947 and 1.1045 respectively. These GC factors are used to removed the effect caused by shared individuals in tissues that may inflate the false-positive rate. To compute a GC factor, one converts the median of the observed p-values into a chi-square statistic, then finds a multiplying factor to scale this new statistic to a chi-square random variable that has p-value at 0.50 (Devlin and Roeder, 1999).

## 3 Results

### 3.1 RECOV controls false-positive rate

We measure the false positive rate (FPR) of RECOV and RE2 at a single variant. To do this, we simulate the data under the null hypothesis where this variant is not associated with the gene expression in any of 44 tissues. To make the data more realistic, we let tissues share individuals where the amount of sharing is the same as in the GTEx data. Moreover, we make expression levels of the same gene from the same individual be correlated with the average correlation of 0.5 across tissues, using the sampling method described in (Sul et al., 2013). This correlation of expression can occur when the tissues of an individual have been exposed to the same environmental factors.

We repeat the simulation 1,000 times at a single variant. In each time, we estimate the effect size and variance of the variant on the gene expression in each tissue. RECOV and RE2 take these effect sizes and variances and produce a meta-analysis p-value for this variant. The GC factor that we estimated previously in section 2.4.2 is applied to this p-value in each simulation. This removes the effect of shared individuals, which is not explicitly modeled in RECOV and RE2. The FPR of this variant is the fraction of times its transformed p-values are significant. We repeat this experiment for 1,000 independent variants, so that we have 1,000 measures of FPR for RECOV and RE2. We use the significance level of 0.05 (*α* = 0.05), and hence we expect FPR of methods to be 0.05 if they control false positives.

We find that RECOV attains correct FPR for the majority of variants that we tested. In RECOV, the median FPR among the 1,000 variants is 0.05 and the 75% and 95% quantiles are 0.06 and 0.09. In RE2, the median FPR is 0.05 and the 75% and 95% quantiles are 0.07 and 0.10. These results demonstrate that both our method and RE2 control the false positive rates with the GC factors that remove the inflation caused by collecting multiple tissues from same individuals.

### 3.2 RECOV discovers more eGenes in GTEx data

We apply our approach to the real multi-tissue eQTL dataset from GTEx. We use GTEx Pilot Dataset V6 released on January 12, 2015. GTEx has performed RNA-seq on 44 tissues from hundreds of individuals, and we select 15,336 genes that have expression data in all 44 tissues. For genotype data, we use GTEx data with imputation that has 5 million genetic variants. For each gene, we use cis-variants that are variants located within 1Mb from its transcription start site. Not all variants are genotyped in every tissue, because the 44 tissues contain samples from different individuals. We use only variants that are genotyped in all 44 tissues. The median number of cis-variants tested per gene is 3,744.

We apply RECOV, RE2, and TBT to the dataset, and for this dataset, we compute eGene p-values for each gene, which is a p-value at a gene level. Before computing an eGene p-value, we apply the GC factor to the observed eGene statistic which is the minimum meta-analysis p-value from all the cis-variants. The GC factor removes the inflation of statistics caused by shared individuals in the tissues from our meta-analysis result. Given all eGene p-values, we use Bonferroni correction to control for multiple testing correction at 5% level to identify significant eGene p-values; thus each gene has a significance threshold of 0.05/15,336.

We find that RECOV detects the most number of eGenes among the three methods from the GTEx dataset. Out of the 15,336 genes tested, RECOV finds that 81.40% of those genes are eGenes while TBT and RE2 find 61.90% and 78.45% of genes are eGenes, respectively. This shows that our approach detects 3% more eGenes than RE2 and about 20% more eGenes than TBT. RECOV detects more eGenes because there are many related tissues in the GTEx dataset. For example, the effects of cis-variants of a gene expressed in brain tissues can be strongly correlated (Figure 4A). It is important to note that a majority of genes are eGenes according to our approach, and this is expected because those are genes in which there is at least one eQTL in at least one of the 44 tissues. Because there are many tissues tested, it is likely that at least one tissue contains an eQTL, which significantly increases the number of eGenes. This result demonstrates that our approach has greater power to detect eGenes in real multi-tissue eQTL datasets than the previous approaches.

**Figure 4:**
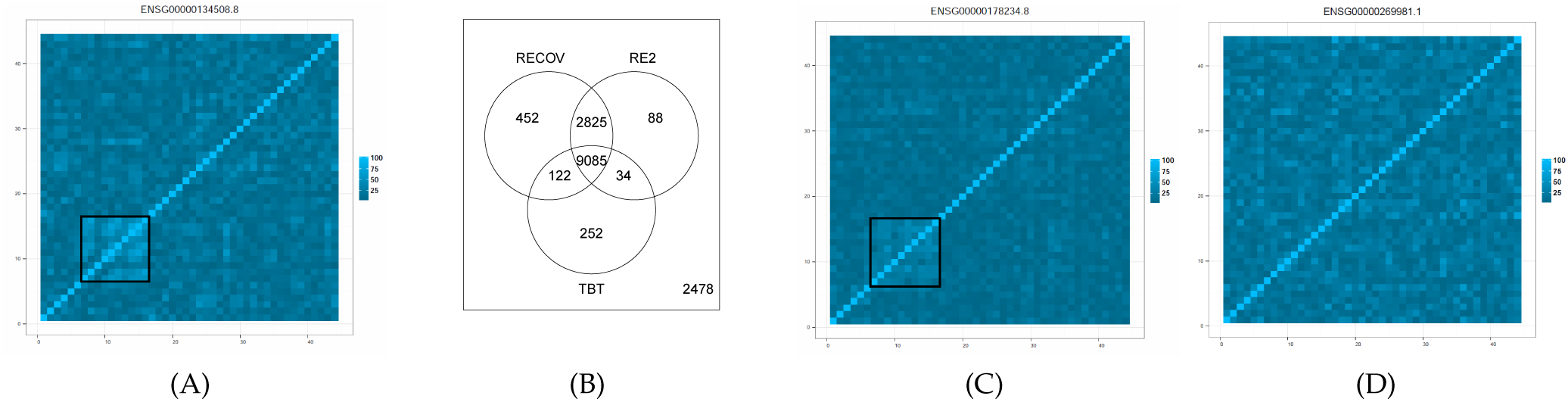
(A) The correlation of SNP-effects for the gene ENSG00000134508.8 in 44 tissues (tissue names are omitted). The correlation is computed by using the matrix *B_g_* in section 2.3.2 where the formula is 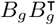 (after proper scaling). Black box indicates the brain tissues. ENSG00000134508.8 is found to be an eGene by only the RECOV method. (B) Venn diagram of the numbers of eGenes found by TBT, RE2, and RECOV. The correlation of SNP-effects for gene (C) ENSG00000269981.1 and (D) ENSG00000178234.8 in 44 tissues (tissue names are omitted). ENSG00000269981.1 and ENSG00000178234.8 are found to be eGenes by only the RE2 method.

Figure 4B shows the Venn diagram of the numbers of eGenes found by TBT, RE2, and RECOV. There are 252 genes detected only in the TBT method. In the TBT method, one analyzes each tissue independently, and is able to determine the tissues in which a gene is an eGene. Thus, TBT has been reported to be the most powerful option to detect genes with eQTLs that have an effect in only one tissue (The GTEx Consortium, 2015). In fact, out of these 252 genes, 225 are eGenes in only 1 tissue, 25 are in 2 tissues, and only 2 are in 3 tissues.

Of the 452 genes discovered by only RECOV, the average RECOV eGene p-value is 8.52E^−9^ (±1.51E^−8^); whereas the average RE2 eGene p-value is 4.18E^−3^(±2.85E^−2^). To understand why RECOV discovers genes which are not found by TBT and RE2, consider the protein-coding gene CABLES1 (Gencode id ENSG00000134508.8) which is only detected by RECOV. From the GTEx portal, CABLES1 is expressed mostly in brain tissues, yet it does not have any brain-specific eQTLs (The GTEx Consortium, 2015). RECOV is a meta-analysis method that pools samples across tissues to increase signals of eQTLs. Thus, it is better than TBT when the sample size is low so that eQTL signals may be undetected. Unlike RE2, the meta-analysis of RECOV considers correlation of the cis-variants across the tissues; thus RECOV would be better than RE2 if CABLES1 has a consistent correlation pattern. This is indeed the case (Figure 4A). CABLES1’s RECOV and RE2 eGene p-value are 4.94E^−13^ and 5E^−5^, respectively.

Of the 88 genes discovered by only RE2, the average RE2 eGene p-value is 1.15E^−8^ (±1.44E^−8^); whereas the average RECOV eGene p-value is 1.85E^−4^(±2.32E^−4^). We suspect that these 88 genes are genes with eQTLs in multiple tissues. However, due to low sample size, the eQTLs signals may be undetected or do not produce an eGene q-value less than the significance threshold in TBT analysis. As a case study, consider the protein-coding gene GALNT11 (Gencode id ENSG00000178234.8) which is detected by only RE2. Like CABLES1, GALNT11 is expressed mostly in the brain tissues (The GTEx Consortium, 2015). Unlike CABLES1, GALNT11 has eQTL signals in the frontal cortex brain tissue, but these signals produce an eGene q-value of 0.0189 which is higher than the TBT significance threshold. In this case, a meta-analysis approach is more suitable. GALNT11’s cis-variants have correlated effect sizes across the brain tissues, but this pattern does not stand out from the rest of the tissues when compared to that of CABLES1 (Figure 4C). For this reason, GALNT11’s RECOV p-value is more than its RE2 p-value (3.50E^−4^ vs 7.08E^−8^). RECOV may also be worse than RE2 when the cis-variants do not have an obvious pattern of correlation across the 44 tissues. As an example, consider the pseudogene RP11-34P13.16 (Gencode id ENSG00000269981.1) which is not tissue-specific (The GTEx Consortium, 2015). The effect sizes of its cis-variants appear to be randomly correlated (Figure 4D), and its RECOV and RE2 p-value are 1.50E^−4^ and 1.37E^−8^, respectively. Altogether, these attributes may have caused the differences between RECOV and RE2.

## 4 Discussion

In this paper, we proposed a novel meta-analysis method called RECOV that better models multi-tissue eQTL datasets to identify eQTLs and eGenes. Based on the biological insight that genetic variants may have similar regulation of gene expression in related tissues, we explicitly modeled in our meta-analysis the correlation of effects that genetic variants have in multiple tissues. We were able to achieve this by extending the previous RE2 model of meta-analysis that assumes independence of effects in multiple tissues. We showed using simulations that our method correctly controls false positive rate. Our method RECOV also has advantage over more advanced models such as eQTLBma and Meta-Tissues, because it can easily handle 44 tissues in the GTEx data. We applied our approach to the real multi-tissue eQTL datasets from GTEx and found that our approach detects more eGenes than the RE2 and TBT analysis. In this paper, our choice of the covariance term in RECOV is heuristic; however, we have introduced a new meta-analysis model where one can use any covariance structure that fits the data better.

In RECOV, we take into account the fact that a genetic variant may have correlated effects in related tissues due to the similar biological mechanisms of its gene regulation. However, it can also have correlated effects when multiple tissues are collected from the same individuals. This correlation is different from the correlation due to the similar biological mechanisms of eQTLs in multiple tissues, and may cause inflation of test statistics for both our approach and the previous RE2 approach. To address this inflation, we measured the genomic control inflation factor from the observed statistics, and this is one of popular approaches to reduce the inflation of test statistics in GWAS. Although we showed that the genomic control approach controls the false positive rate, it may be a conservative approach bearing sub-optimal power. Ideally, one could use the linear mixed models (LMM) approach in Meta-Tissue, which considers correlation due to collection of multiple tissues from same individuals, to generate test statistics that remove this correlation. However, LMM is computationally extensive and requires optimization with the latest techniques (Kang et al., 2010; Zhou and Stephens, 2012). A second option is to directly model this phenomena using the current RECOV approach. One can introduce another random effect corresponding to the fact that tissues may share individuals into the RECOV model. Both of these options are the future direction of our work.

